# MegaBayesianAlphabet: Mega-scale Bayesian Regression methods for genome-wide prediction and association studies with thousands of traits

**DOI:** 10.1101/2022.05.06.490983

**Authors:** Jiayi Qu, Daniel Runcie, Hao Cheng

## Abstract

Large-scale phenotype data are expected to increase the accuracy of genome-wide prediction and the power of genome-wide association analyses. However, genomic analyses of high-dimensional, highly correlated data are challenging. We developed MegaBayesianAlphabet to simultaneously analyze genetic variants underlying thousands of traits using the flexible priors of the Bayesian Alphabet family. As a demonstration, we implemented the BayesC prior in the R package MegaLMM and applied it to both simulated and real data sets. Our analyses show that the resulting model MegaBayesC can effectively use high-dimensional phenotypic data to improve the accuracy of genetic value prediction, the reliability of marker discovery, and the accuracy of marker effect size estimation in genome-wide analyses.

## Introduction

The advent of high density genome-wide single nucleotide polymorphism (SNP) arrays in the past decades has provided exciting new material for the genetic analysis of complex traits. Linear mixed models that can integrate such large-scale genomic data are widely used for genomic prediction (Meuwissen *et al*. 2001; VanRaden 2008) and genome-wide association studies (Visscher *et al.* 2017). Recent advance in multi-omics methodologies now provide opportunities to generate large-scale transcriptomic, metabolomic, and epigenomic profiles as well. The integration of these high-dimensional phenotypes into association studies can increase power to detect causal variants. For example, gene expression profiling in thousands of genes has been used for the identification of genes that affect transcriptional variation (i.e., eQTLs) (Gibson and Weir 2005; McGraw *et al*. 2011), and integrative approaches combining genomic and gene expression data can have higher power to capture the true pathway associations underlying human diseases and complex traits (Xiong *et al.* 2012). In addition, recent developments of highthroughput phenotyping platforms have made the collection of thousands to millions of physiological measurements affordable to breeders (Araus *et al*. 2018). For example, images collected through thermal and hyperspectral cameras are used to increase the accuracy in genomic prediction for grain yield in wheat (Rutkoski *et al*. 2016). To further improve genomic prediction and to understand the underlying genetic mechanism, statistical models that enable the joint analysis of high-dimensional traits are required to establish the connection between phenomics and genomics.

Genomic analyses of high-dimensional, highly correlated data present analytic and computational challenges. The multivariate linear mixed model (MvLMM) is a widely-used statistical model for the genetic analyses of two or more correlated traits (Henderson and Quaas 1976). However, most algorithms used to fit MvLMMs require repeated inversions of genetic and residual covariance matrices among all traits, with a computational burden that grows cubically to quintically as the number of traits increases (Zhou and Stephens 2014). MvLMMs are also susceptible to over-fitting unless sample sizes are very large. Re-parameterizing MvLMMs as Bayesian sparse factor models can alleviate much of this computational burden (Runcie and Mukherjee 2013; Runcie *et al*. 2021) and can significantly improve the accuracy of genomic prediction (Runcie *et al*. 2021). For example, `BSFG` and `MegaLMM` are based on the assumption that the covariances among large sets of traits can be explained by a small set of latent factors (e.g., through gene regulatory networks), which is consistent with the discovery that variation in gene expressions of human diseases are mainly regulated by a few major disease-associated pathways (Xiong *et al.* 2012).

While `MegaLMM` addressed the statistical and computational challenges of applying MvLMMs to high-dimensional phenotypes, it permits a limited range of models for high-dimensional genotype data. Specifically, `MegaLMM` incorporates genomic data through one (or more) genomic relationship matrices, which imposes specific assumptions about the distribution and effect sizes of the underlying genetic variants, and does not allow direct inference on the identities of causal loci. Whole-genome regression methods such as the Bayesian Alphabet methods (Meuwissen *et al.* 2001; Park and Casella 2008; Kizilkaya *et al.* 2010; Habier *et al.* 2011; Cheng *et al.* 2015; Erbe *et al.* 2012; Cheng *et al.* 2018b), on the other hand, encode a wide range of different and more flexible distributions on the effect sizes of causal genomic loci and allow for inference of the causal loci themselves. However, fitting Bayesian Alphabet methods to very large numbers of markers can also be computationally demanding even for a single trait, and extensions of these methods to multivariate traits are very limited.

In this paper, we incorporate whole-genome regression approaches into a Bayesian sparse factor model named *MegaBayesianAlphabet* to incorporate thousands of traits for genome-wide prediction and association studies. The Bayesian Alphabet methods with mixture priors on marker effects (Kizilkaya *et al.* 2010; Habier *et al.* 2011; Moser *et al.* 2015; Wolc *et al.* 2016; Mehrban *et al.* 2017; Wang *et al.* 2020) are popular genetic models due to their incorporation of biologically meaningful assumptions and the variable selection procedure performed during model fitting. We focus on BayesC as an example of a Bayesian Alphabet method (Kizilkaya *et al.* 2010; Habier *et al*. 2011; Cheng *et al.* 2018b), but extensions of MegaBayesianAlphabet with other priors should be straightforward. We show that MegaBayesianAlphabet with BayesC prior (hereinafter referred to as MegaBayesC) can improve genomic prediction accuracy relative to multi-trait GBLUP and RR-BLUP methods by leveraging mixture priors on marker effects and information from thousands of traits. In association studies with millions of markers, MegaBayesianAlphabet is still computationally demanding, but we propose a two-step approach that can accurately estimate marker effects and improve power for association inference in both simulated and real data studies. MegaBayesianAlphabet is implemented in an R package called “MegaLMM”.

## Materials and Methods

In a conventional MvLMM, the genetic and non-genetic correlations among *t* traits are modeled through one or more *t ×t* genetic covariance matrices (**G**_*m*_) and a *t ×t* residual covariance matrix (**R**), respectively. The computational cost of fitting a MvLMM can be prohibitive when *t* is large due to the difficulty in taking inverses of the covariance matrices (Gilmour *et al.* 1995; Yang *et al.* 2011; Zhou and Stephens 2014). To overcome the computational challenge and overfitting in conventional MvLMMs, we reparameterized the conventional MvLMM as a factor model (i.e., MegaLMM (Runcie *et al.* 2021) and MegaBayesianAlphabet), where *K* independent (unobserved) latent factors are introduced to account for the covariances among the *t* traits.

### Model Description

In MegaBayesianAlphabet, the variation among *t* observed traits is decomposed into two parts: the variation caused by dependencies on *K* independent latent factors, which induces correlations among the *t* observed traits, and the variation that is unique, or idiosyncratic, to each trait. In MegaBayesianAlphabet, genetic values of latent factors are defined as a linear combination of all marker effects, and priors from the Bayesian Alphabet methods (Meuwissen *et al*. 2001; Park and Casella 2008; Kizilkaya *et al*. 2010; Habier *et al*. 2011; Cheng *et al*. 2015; Erbe *et al*. 2012; Cheng *et al*. 2018b) are assigned to the marker effects. The model specification of MegaBayesianAlphabet is described below.

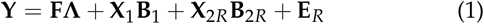

with

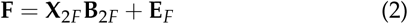

where **Y** is an *n×t* matrix of observations for *n* individuals on *t* traits, **F** is an *n ×K* matrix of latent factors for *n* individuals across *K* latent factors, and **Λ** is a *K × t* factor loading matrix whose elements, such as *λ*_*kj*_, describe the corresponding factor-trait relationships (e.g., the relationship between factor *k* and trait *j*). The *K* latent factors in **F** are further decomposed into genetic effects (i.e., **X**_2*F*_ **B**_2*F*_) and residual effects (i.e., **E**_*F*_) as shown in Equation 2. The genetic effects of latent factors are expressed as multiple regressions on genotype covariates, where **X**_2*F*_ is an *n× b*_2*F*_ matrix of genotype covariates, and **B**_2*F*_ is a *b*_2*F*_ *×K* matrix of marker effects for the *K* latent factors at *b*_2*F*_ genotyped loci. **X**_1_ is an *n × b*_1_ incidence matrix allocating the observations on *t* traits to *b*_1_ fixed effects with coefficient matrix **B**_1_. The residuals are similarly decomposed into trait-specific genetic effects (i.e., **X**_2*R*_ **B**_2*R*_) and trait-specific residual effects (i.e., **E**_*R*_), with **B**_2*R*_ being a *b*_2*R*_ *×t* matrix of marker effects corresponding to the *t* traits at *b*_2*R*_ genotyped loci.

If all sources of correlation among observed traits are explained by the latent factors, the residuals conditional on these factors become uncorrelated between different traits. Since the sources of correlation among observed traits are explained by independent latent factors, samples at each iteration of Markov chain Monte Carlo (MCMC) can be obtained simultaneously in parallel across traits and factors, which leads to significant reduction in the computational cost of model fitting.

### Prior Specification

#### Genetic Marker Effects

Mixture priors are widely used for genetic marker effects in Bayesian regression methods in genome-enabled analysis. In this paper, the BayesC prior is used for the marker effects (e.g., coefficients in **B**_2*F*_) and we term this specific version of MegaBayesianAlphabet: MegaBayesC. The BayesC mixture prior assumes that marker effects are independently and identically distributed, each of which has a point mass at zero with a marker exclusion probability *π*, and follows a univariate normal distribution with a marker inclusion probability 1 *−π*. For example, the prior distribution of the marker effect at locus *i* for the *k*th latent factor is shown as follows.

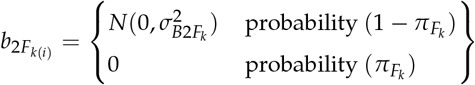

where 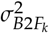 is the variance of marker effects corresponding to factor *k*. Due to the independence among latent factors, and the independence among traits conditional on **FΛ**, marker effects can be efficiently sampled from a set of univariate BayesC models in parallel across traits and factors at each iteration of MCMC. We treat each marker exclusion probability for the *K* latent factors (e.g.,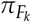) and the *t* observed traits as an independent unknown parameter to be estimated. Note that if marker inclusion probabilities for all factors were set to 1.0 (i.e., all markers are included with equal variance), the model is equivalent to RR-BLUP, and we term this specific version of MegaBayesianAlphabet: MegaRRBLUP.

#### Factor Loading Matrix

The factor loading matrix (**Λ**) describes the relationship between latent factors and observed traits. Sparsity in this matrix implies that factors affect some, but not all traits, a key assumption in Bayesian sparse factor models (Carvalho *et al.* 2008). We use a BayesC mixture prior for the elements of **Λ**. Because factor swaps do not change the likelihood, to improve the identifiablilty of the model, we introduce an additional parameter to the included-variable variance 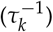 that is stochastically decreasing across factors (Bhattacharya and Dunson 2011; Runcie and Mukherjee 2013; Runcie *et al.* 2021). For the factor loading that describes the relationship between factor *k* and trait *j* (i.e., *λ*_*kj*_), its prior distribution is shown as follows.

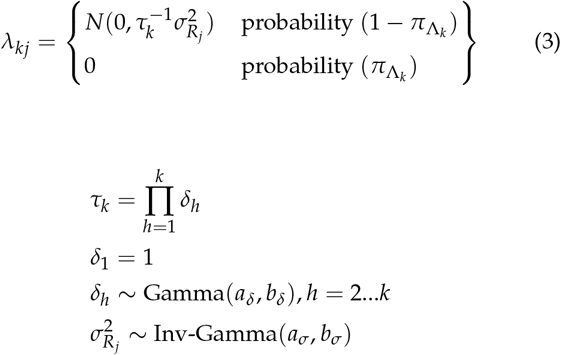

Through the prior specification of **Λ**, an appropriate level of truncation on the rows of **Λ** is able to ensure that the contribution from additional factors beyond the truncation point is negligible (Bhattacharya and Dunson 2011).

#### Other priors

All other prior distributions are the same as used in Runcie *et al*. (2021).

#### Posterior Distributions for Gibbs Sampler

We use MCMC method to sample from the posterior distributions of all parameters. The full conditional posterior distributions for Gibbs sampler are derived for all the parameters in MegaBayesC in **Appendix**.

#### Estimation of Genetic Values for Genomic Prediction

We assessed the performance of MegaBayesC as a tool for genomic prediction using hyperspectral data as additional traits to assist wheat yield prediction.

#### Data Description

Best linear unbiased estimators (BLUEs) of grain yield and reflectances from 62 wavelength bands collected with an areal hyperspectral camera on each of 10 time-points during the growing season for 1033 bread wheat lines were downloaded from CIMMYT Research Data (Krause *et al.* 2019). We analyzed results from the 2014-2015 breeding cycle under the Optimal Flat treatment. All lines were genotyped using the pipeline described in Poland *et al*. (2012). Markers with call rate ≤50% and minor allele frequency (MAF) ≤ 0.05 were removed. Missing genotypes were imputed by corresponding marker means. In our analysis, the 620 hyperspectral BLUEs were used as secondary traits (Runcie and Cheng 2019) to improve the prediction of the genetic value of grain yield, which is served as a focal trait in our prediction scenario. Both sets of traits were combined into a 1033 *×* 621 trait matrix **Y**.

#### Models

Four different models were used to predict the grain yield (GY): GBLUP, MegaGBLUP, MegaRRBLUP, and MegaBayesC. Posterior means were used as point estimates of parameters of interest. These four models are described below. **GBLUP** A conventional single-trait GBLUP model (VanRaden 2008) with a variance-covariance matrix proportional to a genomic relationship matrix **K** fitted to the grain yield BLUEs, ignoring the hyperspectral data.

### MegaGBLUP

This model was described in Runcie *et al*. (2021). The fixed effects **B**_1_ included intercepts only. The random effects **B**_2*R*_ and **B**_2*F*_ were not included in the model. A random effect with covariance proportional to **K** was included in Equations 1 and 2 to model the genetic relationships among lines.

### MegaBayesC

The estimated individual genetic merits of grain yield in MegaBayesC were computed as: 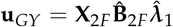, where 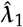 denotes the first column of 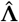, which specifies the estimated relationship between all factors and grain yield. **B**_1_ included only an intercept, and **B**_2*R*_ was not included. We included one factor having non-zero effects only on grain yield, i.e., 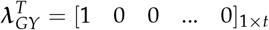, to model direct genetic effects on grain yield. For the remaining factors, the probability of a element from **Λ** being zero was considered as known and set to be 0.9 to introduce sparsity to **Λ** and to shorten the time required for its convergence, while the probability of a marker having a null effect on a latent factor was considered as unknown and was estimated. *K* = 100 factors were fitted in MegaBayesC.

### MegaRRBLUP

This model mimics the priors for marker effects in RR-BLUP. The only difference between MegaRRBLUP and MegaBayesC lies in the prior distributions of marker effects on latent factors. Normal distributions instead of mixture priors are used for marker effects in MegaRRBLUP, indicating that all markers are included in the model. This model should be identical to **MegaGBLUP** except that the prior on elements of **Λ** is BayesC instead of the Bayesian Horseshoe.

#### Cross Validation

We used cross-validation to evaluate the predictive performance of different models by masking the grain yield of 516 randomly selected lines (around 50% of the population) of the population during model fitting and comparing the masked values to model predictions. Since we did not mask the hyperspectral data from these 516 model validation individuals, but used those data to enhance our genetic value predictions, using the Pearson’s correlation between predicted and observed GY values could lead to biased and sub-optimal choices of models (Runcie and Cheng 2019). Instead, we used the estimated genetic correlation corrected by grain yield heritability to estimate the prediction accuracy (Runcie and Cheng 2019; Daetwyler *et al*. 2013) as implemented in Runcie *et al*. (2021). The cross-validation process was repeated 20 times with different masked lines.

#### Estimation of Marker Effects for Association Inference

In MegaBayesianAlphabet, covariances among high-dimensional phenotypic data are decomposed into *K* sources of variation, each of which controls the correlation among a subset of observed traits through the factor loading matrix. In this way, information of correlated traits is used jointly to estimate their underlying pathways (i.e., latent factors), while the computational burden to analyze large-scale phenotypic data is significantly decreased. With the assistance of large-scale genetically correlated traits, MegaBayesianAlphabet is expected to boost the discovery of genetic variants associated with a trait of most interest (i.e., focal trait) and precisely quantify their effect sizes.

In this section, two simulation studies and one real data analysis were conducted to investigate the accuracy of the estimation of marker effects by MegaBayesC. First, a population with independent and uncorrelated SNPs was simulated to demonstrate the ability of MegaBayesC to distinguish the genetic and non-genetic sources of variation in a focal trait, utilizing the information of correlated traits. Second, a simulation study based on a real Arabidopsis population was conducted to study the effects of population structure and linkage among markers. Finally, the real phenotypes of flowering time from this Arabidopsis population was studied, utilizing expression data from 20,843 genes.

#### Simulated Study in A Population without structure or Linkage Disequilibrium

We created a simulated population of *n* = 5000 individuals and *p* = 2, 000 SNPs. An *n×p* matrix of genotypic covariates was generated by random sampling from {0, 1, 2}. We then created simulated phenotypic data for a single focal trait and many correlated “secondary” traits. The performance of MegaBayesC was compared to a single-trait BayesC model (ST-BayesC) based on the accuracy of estimated marker effects for the focal trait. We induced genetic and non-genetic covariation among the traits through latent factors. The majority of variance in the focal trait was attributed to the latent factors. In Scenario 1, we created latent factors whose variation was primarily determined by the genetic markers (i.e. high-heritability latent factors), and in Scenario 2, the latent factors were predominantly non-genetic (i.e. low-heritability factors).

We studied four parameters that we expected to influence the relative performance of MegaBayesC and ST-BayesC. They are 1) the number of latent factors (*n* _*f actor*_), 2) the number of correlated traits (*n*_*trait*/ *f actor*_) controlled by each factor, 3) the number of QTL (i.e., causal variants) that control each factor (*n*_*qtl*/ *f actor*_), and 4) the heritability of the factors. In this simulation study, *n* _*f actor*_ = *{*2, 6, 9*}, n*_*trait*/ *f actor*_ = *{*2, 20*}, n*_*qtl*/ *f actor*_ = {10, 20, 30}, and two heritable patterns of latent factors were considered.

To generate the simulated phenotype data, we first used *n* _*f actor*_ and *n*_*trait*/ *f actor*_ to construct a factor loading matrix (**Λ**). For example, when *n*_*trait*/ *f actor*_ = 2 and *n* _*f actor*_ = 2, 4 (i.e., *n* _*f actor*_*× n*_*trait*/ *f actor*_) observed traits were simulated, with two different observed traits linked to each factor. Since the first observed trait was treated as the focal trait, and all factors were assumed to contribute to its variation, factor loadings in the first column of **Λ** were set to 1. To minimize the complexity of this simulation, non-zero factor loadings in **Λ** were set to be equal to 1. Therefore, the simulated **Λ** given *n*_*trait*/ *f actor*_ = 2 and *n* _*f actor*_ = 2 was expressed as:

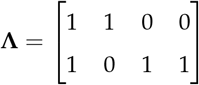

Based on the constructed **Λ**, all factors except the first factor were linked to *n*_*trait*/ *f actor*_ + 1 = 3 observed traits, while the first factor was linked to *n*_*trait*/ *f actor*_ = 2 observed traits. A similarly structured **Λ** was constructed for other combinations of *n*_*trait*/ *f actor*_ and *n* _*f actor*_.

After defining **Λ**, genetic variation controlled by selected QTL and non-genetic variation in each factor were simulated. *n*_*qtl*/ *f actor*_ QTL were selected for each factor, and variation was simulated such that the variance explained by these QTL was a defined percentage of the total variation in the factor. In Scenario 1, the QTL accounted for 95% of the variance of each factor (i.e., 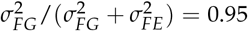 with 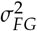 being the genetic variance of factors and 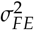 being the residual variance of factors). In Scenario 2, only the first factor was associated with QTL (again with 95% of its variance explained by the QTL), and the remaining factors had independent variation. Finally, additional trait-specific variance was added to each trait, accounting for approximately 10% of its total variance.

As a consequence of these simulation choices, the two scenarios differed in several key aspects of the genetic architecture and correlation structures between the focal trait and the secondary traits. In Scenario 1, all factors were controlled predominately by genetic variation and all QTL for every factor was therefore a QTL for the focal trait. Therefore, all secondary traits had strong genetic correlations with the focal trait. In Scenario 2, most factors were controlled by non-genetic variation; only the first factor was controlled by QTL. Therefore while all secondary traits were phenotypically correlated with the focal trait, most of these correlations were non-genetic.

In both scenarios, as *n* _*f*_ *actor* and/or *n*_*qtl*/ *f actor*_ increased, the magnitude of variation explained by each QTL decreased to hold the total percentage of variation in the focal trait explained by QTL constant. In Scenario 1, when *n* _*f actor*_ = 9 and *n*_*qtl*/ *f actor*_ = 10, the 90 QTL each explained ≈ 0.97% of the total variance (Figure 3). As *n*_*qtl*/ *f actor*_ increased to 30, the number of QTL for the focal trait increased to 270 and each accounted for around 0.29% of the total variance of focal trait. In Scenario 2, when *n* _*f actor*_ = 9, the variance explained by each marker decreased from 0.90% to 0.31% as the number of QTL increased from 10 to 30. For a given *n* _*f actor*_ and *n*_*qtl*/ *f actor*_ the per-QTL effect sizes were comparable, but since there were more factors with QTL in Scenario 1, the total variance in the focal trait controlled by all QTL was larger.

In Scenario 2, as *n* _*f actor*_ increased, the proportion of variance explained by QTL decreased. For example, when *n* _*f actor*_ = 6, the genetic variance accounted for 14% of the total variance of focal trait. With *n* _*f actor*_ = 9, the percent of variance explained by genetic markers decreased to 9%. In Scenario 1, the percentage of variance explained by QTL was constant across values of *n* _*f actor*_. In this scenario, all QTL for all factors contributed to the variation in the focal trait. For example, when *n* _*f actor*_ = 6 and *n*_*qtl*/ *f actor*_ = 10, each factor was influenced by 10 QTL, which were randomly selected from all SNPs, leading to a total of 60 QTL selected. During this process, some SNPs may be stochastically selected more than once, and thus, some QTL may have effects on more than one factor.

Based on the combination of *n*_*trait*/ *f actor*_, *n* _*f actor*_, *n*_*qtl*/ *f actor*_, and the heritable patterns, a total of 3 *×* 3 *×* 2 *×* 2 conditions were studied in this simulation study. 10 replicates were conducted for each of the 36 conditions.

When fitting models to these simulated data, we included the intercept for each trait as the only fixed effect. The model specification of MegaBayesC was similar to that used in the **Genomic Prediction** application, except: 1) no fixed factor loadings were included in **Λ**, 2) the probability of a element from **Λ** being zero was considered as unknown and was estimated, 3) the number of factors fitted in the model was *K* = 10. We estimated the total marker effects on the focal trait obtained by MegaBayesC as: ***α*** _*f*_ = **B**_2*F*_ ***λ***_1_, where **B**_2*F*_ is the matrix of marker effects of latent factors and ***λ***_1_ denotes the first column of **Λ** specifying the relationship between factors and focal trait, i.e., summing up the QTL effects on the latent factors weighted by the relationships between each factor and the focal trait. We measured the performance of each method (i.e., MegaBayesC and ST-BayesC) by calculating the square root of the mean square error (RMSE) of estimated marker effects.

#### Simulation Study in a Real Arabidopsis Population

We created a second set of simulated datasets based on real genotypes from 1003 Arabidopsis thaliana accessions. Genotype data were downloaded from the 1001 genomes project (Alonso-Blanco *et al*. 2016). In a real population, the presence of linkage disequilibrium (LD) between loci and variable allele frequencies among markers increase the complexity of genetic association analyses. We removed SNPs with MAF ≤0.05 and missing genotype rate ≥ 0.1 using PLINK 1.9 (Purcell *et al.* 2007), leaving 802,427 variants used for downstream analysis.

To ensure the QTL were independent, we pruned SNPs with an LD threshold of 0.8 in windows of 500 SNPs, using a sliding window of 100 SNPs. We randomly selected 20 QTL from these SNPs, and generated 10 latent factors, each was affected by 2 different QTL. In this simulation, the structure of the variance of the focal trait was simplified. All genetic variance in all traits was driven by the QTL effects on the latent factors, while all non-genetic variance was trait-specific. In this way, the observed traits (**Y**) was expressed as: **Y** = **X**_2_ **B**_2*F*_ **Λ** + **E**_*R*_.

We set each element of the first column of **Λ** to 0.5 so that all 10 of the factors contributed equally to the focal trait. Each factor was additionally linked to 20 different secondary traits with factor loadings equal to 1. Other elements in **Λ** were set to be 0. Therefore, a total of 201 traits were simulated.

The proportion of genetic variance in the focal trait was set to be around 60% (i.e., 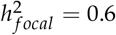). To ensure that the variance explained by each QTL was consistent (≈1 *−* 5% of the total variance), QTL effects were sampled from a uniform distribution *U*(3, 5), and a randomly chosen half of those effects were multiplied by −1. In addition, since the heritability of secondary traits such as gene expression is often higher than that of focal trait in real data applications, the heritabilities of the 200 secondary traits were each set to be 0.8.

Finally, to parallel our real data analysis below, secondary trait data was simulated for only 649 of the 1003 Arabidopsis accessions. The 354 remaining accessions had the records for only the focal trait.

After creating the simulated data, we applied three methods to identify QTL and estimate their effects on the focal trait: 1) single-trait Genome-Wide Association Studies (GWAS) using GCTA (Yang *et al.* 2011) (ST-GCTA); 2) single-trait BayesC implemented in JWAS (Cheng *et al.* 2018a) (ST-BayesC); and 3) MegaBayesC implemented in MegaLMM. Since whole-genome regression models with hundreds of thousands of candidate markers are computationally prohibitive, a two-stage analysis was implemented for ST-BayesC and MegaBayesC. In the first stage (i.e., the pre-selection stage), we selected a small proportion of SNPs to take forward into a full BayesC analysis by running a single-trait GWAS using GCTA on only the 354 individuals without records on secondary traits. After running the GWAS, we used LD-based clumping to select ≈ 2000 potentially important SNPs. First, we sorted SNPs by p-value, removed SNPs with p-values larger than 0.01, then used a greedy algorithm to select the most-significant SNPs and mask all nearby SNPs (within 250Kb) with *r*_2_ *>* 0.5 (Purcell *et al*. 2007).

After the pre-selection stage, the records of focal and secondary traits from the remaining 649 individuals, which are considered as an independent population, were analysed in MegaBayesC using only the pre-selected potentially important SNPs. The model specification of MegaBayesC was similar to that used in the previous simulation study for independent population, except we set *K* = 30. In MegaBayesC, the total marker effects of the focal trait were computed as: ***α*** _*f*_ = **B**_2*F*_ ***λ*** _*f*_, where **B**_2*F*_ is a *b*_2*F*_ *×K* matrix of marker effects for latent factors, with *b*_2*F*_ being the number of SNPs selected at the pre-selection stage, and ***λ*** _*f*_ denotes the column of **Λ** that specifies the relationship between factors and focal trait. Furthermore, to demonstrate that the improved performance of MegaBayesC is attributed to not only the use of the BayesC prior on the marker effects but also the utilization of information from correlated secondary traits, a ST-BayesC was also performed for the 649 individuals at the second stage. MCMC chains of 50,000 iterations were run for the BayesC-based methods with the first 10,000 iterations discarded as burn-in.

In addition to the two-stage analysis, a one-stage ST-GCTA was performed using the whole-genome SNP information and the phenotypes of the focal trait from all 1003 individuals. To compare with the two-stage analysis, the selection of potentially important SNPs was done based on the one-stage ST-GCTA result in the same manner as that in the pre-selection stage.

Figure 1 shows the procedures performed to estimate marker effects in the three different methods. Simulations were repeated 100 times.

**Figure 1.**
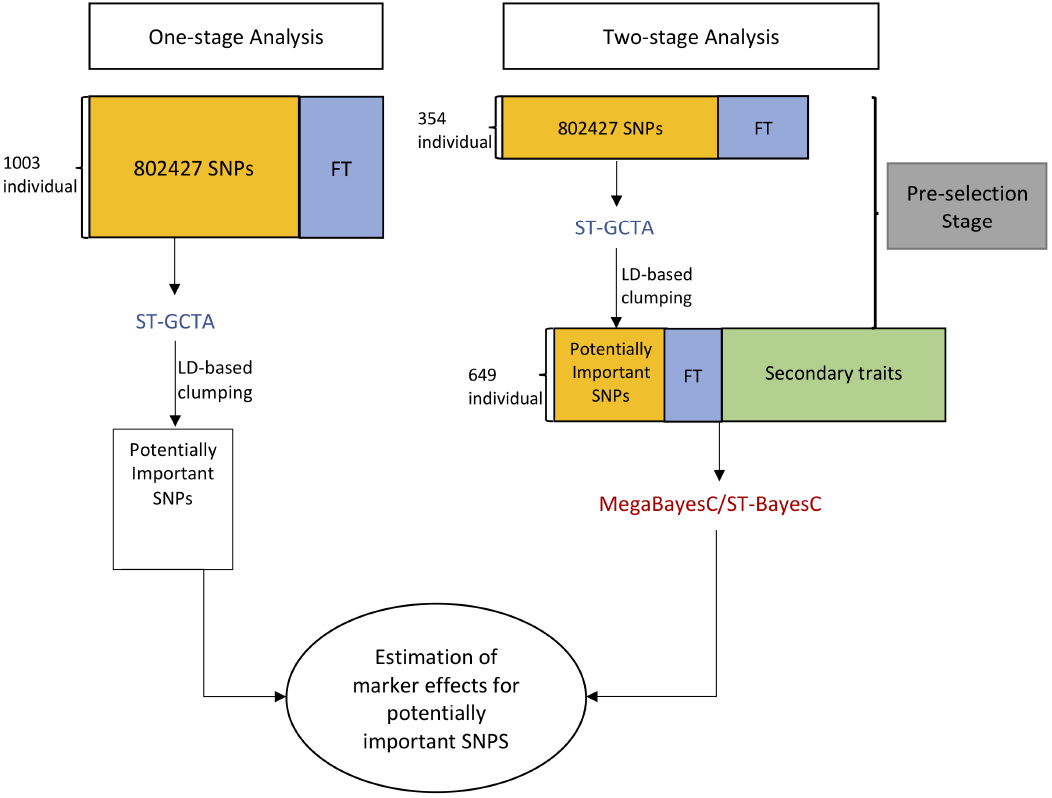
Graphic representation for the procedure of one-stage and two-stage analyses performed for the estimation of marker effects. FT represents the focal trait, ST-GCTA represents single-trait GWAS implemented in GCTA, and ST-BayesC represents single-trait BayesC method. In ST-BayesC, only phenotypes of FT and genotypes of the pre-selected potentially important SNPs were used.

RMSEs of the QTL effect sizes and the percentage of variance in the focal trait explained by the QTL were used to evaluate the accuracy of estimation of QTL effects by different methods. The variance explained by marker *l* was computed as: *var*(*α*_*l f*_ **x**_2_), where *α*_*l f*_ is the marker effect of SNP *l* on focal trait, and **x**_2_ is the vector of genotypic covariates for SNP *l*. To score QTL accuracy, we parsed the detected QTL in three ways: 1) If the true QTL were selected in the set of potentially important SNPs (e.g., Stage 1), the estimated effects were compared directly to the true effects. 2) If a SNP in imperfect LD with the true QTL was selected instead of the true QTL, we flagged its estimated effect size in the accuracy comparison because the incomplete linkage and different allele frequencies of the two SNPs mean that the estimated effect size will not be directly comparable to that of the true QTL. However, the percentage of variance attributed to the marker should be similar to the true QTL as long as *r*_2_ is high; 3) If neither the true QTL nor any of its linked SNPs was selected in the potentially important SNPs, we set the estimated marker effect and variance explained by this QTL to 0. For the purpose of unit consistency, RMSE of estimated marker effects and estimated marker-explained standard deviation (i.e., square root of marker-explained variance) across different methods were compared. In this study, a SNP was considered to be linked to a QTL if the squared correlation between its genotypic covariate and the QTL genotype was greater than 0.4.

#### Genetic Association Analysis of Arabidopsis thaliana Flowering Time and Gene Expression

Phenotypes of flowering time from 1003 accessions and expression data of 20843 genes from 649 accessions were used, with flowering time selected as our focal trait. Gene expression data was downloaded from NCBI GEO (Barrett *et al.* 2012). Genes with average counts smaller than 10 were removed and the remaining gene counts were normalized and variance stabilized as per Runcie *et al*. (2021) using DESeq2 (Love *et al.* 2014). The two-stage MegaBayesC and one-stage ST-GCTA analyses described above were performed again on this dataset. LD-based clumping was done to select potentially important SNPs for both methods. The model specification of MegaBayesC was similar to that used in the **Genomic Prediction** section above. A MCMC chain of 80,000 was run with the first 20,000 iterations discarded as burn-in. In the two-stage MegaBayesC analysis, potentially important SNPs with explained proportion of variance *>* 0.1% were classified as significant SNPs, while in the one-stage ST-GCTA analysis, potentially important SNPs with p-value *<* 1 *×*10_*−*5_ were classified as significant SNPs. We compared each significant SNP to a list of genes previously known to influence flowering time in Arabidopsis (Bouché *et al.* 2016), and counted as a match (i.e., a true positive hit) if a SNP was within +/-100 Kb distance from at least one of the reported genes. Otherwise the significant SNP was conservatively considered as a false positive association.

#### Data availability

Scripts for running all analyses are archived at GitHub: https://github.com/Jiayi-Qu/Mega-BayesC. The Bayesian Alphabet implementation is available on the “BayesAlphabet” branch of the MegaLMM GitHub repository: https://github.com/deruncie/MegaLMM/tree/BayesAlphabet. Data from the wheat breeding trial were downloaded from CIMMYT Research Data (Krause *et al.* 2019). Arabidopsis flowering time data was downloaded from Arapheno: https://arapheno.1001genomes.org/phenotype/261/. Gene expression data was downloaded from NCBI GEO (Barrett *et al.* 2012). Genotype data were downloaded from the 1001 genomes project (Alonso-Blanco *et al*. 2016).

## Results

### MegaBayesC Improves Estimation of Genetic Values

We tested if MegaBayesianAlphabet models could match or exceed the performance of MegaLMM in trait-assisted genomic prediction using data from a breeding trial of bread wheat. We compared the genomic value prediction accuracy of MegaBayesC and MegaRRBLUP to MegaGBLUP in this dataset, where we leveraged 620 hyperspectral phenotypes measured on 1033 bread wheat lines to supplement genotype-based predictions of genomic value for grain yield. As a baseline, we performed conventional univariate GBLUP-based genomic value prediction as well. Prediction accuracy was assessed by cross-validation where for each of 20 replicates, grain yield values of 50% of the lines were masked and used as an independent testing set. Estimated genetic correlations between predicted and observed yields in the testing set were used as the cross-validation statistic.

As shown in Figure 2, univariate GBLUP achieved a prediction accuracy of 0.43 in this dataset. MegaGBLUP fitted to all traits in MegaLMM with a single random effect based on the genomic relationship matrix **K** achieved an average prediction accuracy of 0.69. MegaRRBLUP fitted in MegaBayesianAlphabet achieved an average prediction accuracy of 0.68. No significant difference was observed between MegaGBLUP and MegaRRBLUP. RR-BLUP and GBLUP are mathematically equivalent (Whittaker *et al*. 2000; Meuwissen *et al*. 2001; Habier *et al*. 2007) models that account for the contributions of the genetic markers, but MegaGBLUP uses a horseshoe prior for the elements of **Λ** while MegaRRBLUP uses the BayesC prior for these parameters with fixed *π* = 0.9. MegaBayesC, with its BayesC prior on the marker effects, achieved an average accuracy of 0.75, significantly higher than the other methods. These results show that the use of biologically meaningful prior on marker effects can further improve the genomic selection in breeding programs.

**Figure 2.**
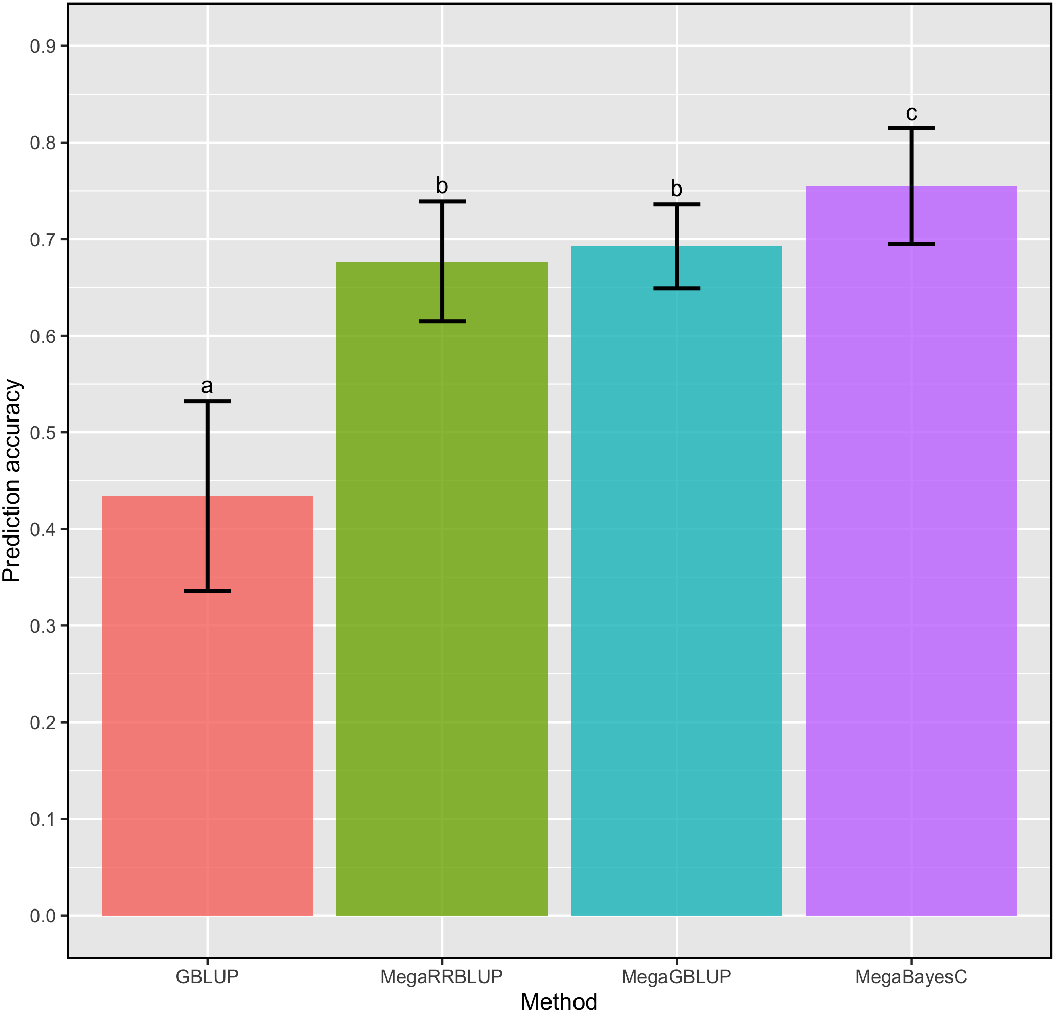
Genomic value prediction performance of 4 models for wheat yield. Records of yield, 620 hyperspectral phenotypes, and genotype data for 1033 lines were available. 20 replicate validations were used. Bars show the mean prediction accuracy (± standard error) for each model, and letters show the statistical significance of mean difference between methods based on a paired t-test.

### MegaBayesC improves Estimation of Marker Effects in Simulated Populations with Independent Markers

Next, we ran a set of simulations to evaluate the ability of MegaBayesC to identify and accurately estimate the effect sizes of genetic variants for a set of correlated traits under different genetic architectures. Specifically, we tested whether MegaBayesC improved the estimation of variant effect sizes of a single focal trait when phenotypes of other correlated traits (i.e., secondary traits) were provided.

Since the magnitude and causes (genetic vs. non-genetic) of the covariance structures among traits determine the usefulness of the secondary traits, we considered two covariance structures. In both cases, we began by simulating a set of latent factors partially controlled by genetic variation. In Scenario 1, the majority of variation in the focal trait was controlled by latent factors that were dominated by genetic variation. In Scenario 2, the majority of variation in the focal trait was controlled by latent factors dominated by non-genetic sources of variation. We compared the estimation of marker effects between ST-BayesC (which ignored all secondary traits) and MegaBayesC (which used all trait data at once). We scored the accuracy of each method by the RMSE of estimated marker effects. In both scenarios, as the genetic architecture increased in complexity (i.e., the number of QTL increased and the average size of each QTL decreased to keep the total percentage of variation attributable to the QTL constant), the performance of ST-BayesC decreased (RMSE increased) much more dramatically than MegaBayesC. Figure 3 shows RMSE of estimated effects for QTL and SNP, respectively, (*i*.*e*. QTL are markers with a non-zero effect and SNPs are markers with a true effect size of zero) under the two scenarios for the simulation setting where the largest difference of RMSE was observed between MegaBayesC and ST-BayesC, with *n*_*trait*/ *f actor*_ = 2 and *n* _*f actor*_ = 9. Results for other combinations of *n* _*f actor*_, *n*_*trait*/ *f actor*_, and *n*_*qtl*/ *f actor*_ are shown in **Appendix** (Figure 9). In Scenario 1, the number of latent factors had no direct effect on the performance of ST-BayesC beyond its effect on the number of QTL. Also, the number of traits linked to each factor (i.e. *n*_*trait*/ *f actor*_) did not significantly affect the performance of MegaBayesC in both Scenario 1 and Scenario 2. This shows the ability of MegaBayesC to capture the underlying sources of correlations among traits by optimizing the utilization of secondary traits, even when each factor only has one linked secondary trait included in the model.

**Figure 3.**
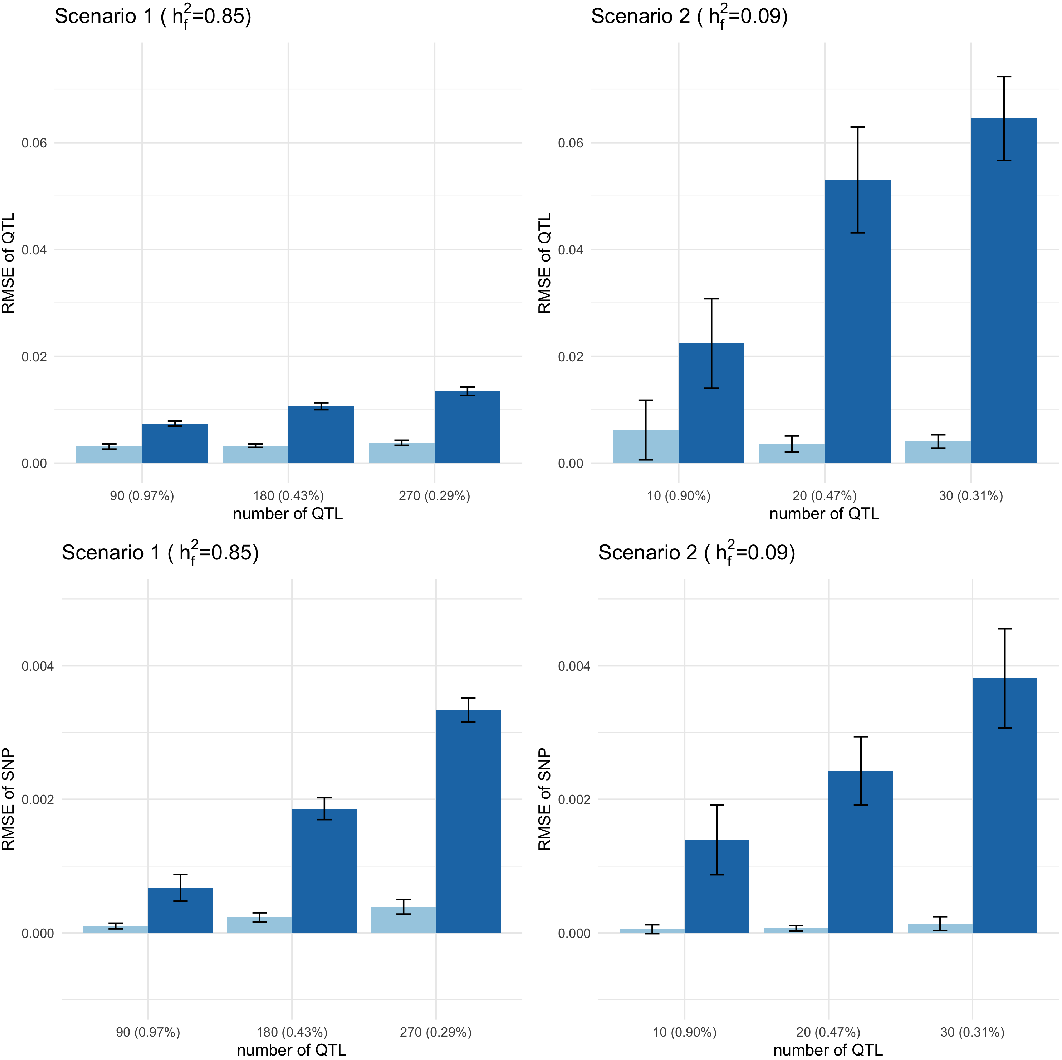
Root mean square error (RMSE) of estimated QTL effects and SNP effects, respectively, under two scenarios.The upper panels show RMSE of estimated QTL effects under two scenarios. The lower panels show RMSE of estimated SNP effects under two scenarios. The left panels show RMSE for Scenario 1, where all latent factors had high heritability (*h*^2^ = 0.95). The right panels show RMSE for Scenario 2, where only one of the factors had high heritability (i.e., factor 1 had *h*^2^ = 0.95 and the remainder factors had *h*^2^ = 0). Results are shown for the simulation setting with *n*_*trait*/ *f actor*_ = 2 and *n* _*f actor*_ = 9. The average proportion of total variance explained by one QTL was shown in the parenthesis.

For ST-BayesC, the RMSE of estimated marker effects increased significantly as marker-explained variances decreased in both scenarios. Compared to Scenario 1, the increase of RMSE for estimated effects of QTL was greater in Scenario 2, while the increase of RMSE for estimated effects of SNPs were similar between the two scenarios. This indicates that the performance of ST-BayesC to identify QTL was affected by the marker-explained variance as well as the variance structure of the focal trait.

In contrast, the performance of MegaBayesC was relatively constant across scenarios as measured by RMSE. In terms of the estimation of effect sizes of QTL, the influence of the variance structure and the marker-explained variance was negligible, which lead to a relatively constant RMSE across the simulation settings. At the same time, MegaBayesC was able to shrink most SNPs more effectively towards zero, especially in Scenario 2, when the ratio of number of QTL to number of SNPs was smaller.

To further explore the differences in the performance of ST-BayesC and MegaBayesC in Scenario 2, we plotted the estimated marker effects under one example simulation with *n* _*f actor*_ = 9, *n*_*qtl*/ *f actor*_ = 30, and *n*_*trait*_ = 2 (Figure 4).

**Figure 4.**
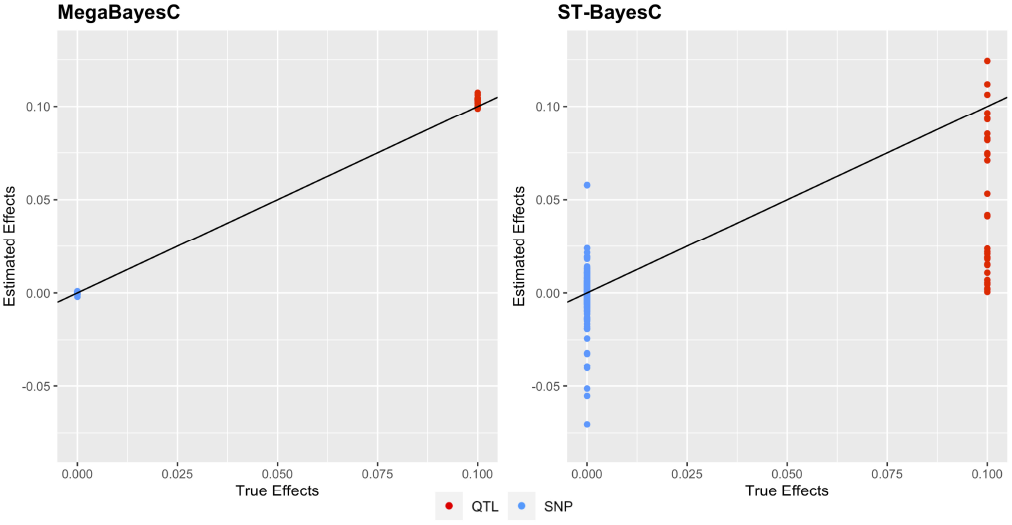
Scatter plot of estimated marker effects versus true marker effects for the simulation setting with *n* _*f actor*_ = 9, *n*_*qtl*/ *f actor*_ = 30, and *n*_*trait*_ = 2 in Scenario 2, where all factors have effects on the focal trait but only one of them is a genetic factor (i.e., *h*^2^ *>* 0). Red and blue colors specify QTL (effect size ≠ 0) and SNP (effect size ≠ 0), respectively. The solid black line represents the line y = x.

For ST-BayesC, some QTL were successfully select by the model and their effect sizes were accurately estimated close to the true value of 0.1. However, for the majority of QTL, the estimated marker effects were shrunk toward 0s. On the other hand, ST-BayesC erroneously estimated effect sizes of SNPs with true effect sizes of 0 from -0.07 to 0.06. In contrast, the marker effects of QTL and null-effect SNPs were accurately estimated by MegaBayesC (Figure 4).

### Estimation of Explained Variance of Markers in a Population Simulated Using Real Genotype Data

To explore the ability of MegaBayesC to accurately identify QTL and estimate their effect sizes in the presence of LD, we generated simulated phenotypes based on real genotypes from an Arabidopsis population. We then ran association analyses using three methods: The direct (*i*.*e*. one-stage) method, ST-GCTA, that only uses the focal trait, and two two-stage methods: ST-BayesC and MegaBayesC, which both rely on a pre-selection stage to select a set of candidate SNPs using one partition of the population, and then an assay stage where the effects of those SNPs on the focal trait are modeled in the second partition of the population. We compared the performance of the models by the RMSE of estimated marker effects and marker-explained variances.

Figure 5 shows the RMSE of estimated marker effects and estimated marker-explained standard deviations from the simulated phenotype data. The two-stage MegaBayesC method achieved the lowest RMSE for both marker effects and marker-explained standard deviations, followed by the two-stage analysis incorporating ST-BayesC, and then the one-stage single-trait GWAS (ST-GCTA). The RMSE of the one-stage single-trait GWAS was around ten times larger than that of the two-stage BayesC-based analyses, while the difference between ST-BayesC and MegaBayesC was much smaller. Furthermore, the RMSE of estimated marker-explained standard deviations was generally lower than that of estimated marker effects. The larger RMSE of estimated marker effects is likely due to the selection of linked SNPs rather than the true causal QTL in the pre-selection stage.

**Figure 5.**
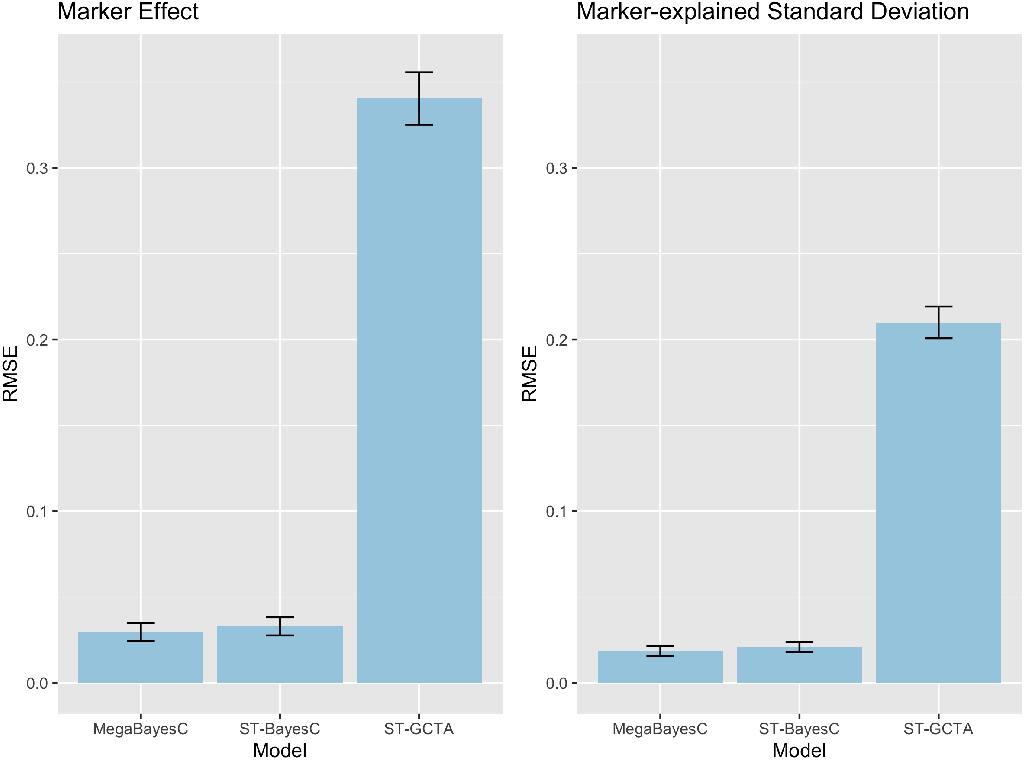
Root mean square error (RMSE) of estimated marker effects and estimated marker-explained standard deviations (i.e., square root of marker-explained variances) across different methods. The performance of two-stage (ST-BayesC and MegaBayesC) methods and one-stage (ST-GCTA) method were compared.

To further explore the difference in the performance of ST-BayesC and MegaBayesC in this simulation scenario, we present the relationship between true and estimated marker effects for one replicate in Figure 6. In this simulation, 19/20 true causal QTL were selected by ST-GCTA, and only 16 were selected in the pre-selection stage for the two-stage methods, ST-BayesC and MegaBayesC. In all these three cases, the effect sizes of these selected QTL were accurately estimated. However, the effect sizes of many null-effect SNPs were dramatically overestimated by ST-GCTA, leading to an overall high false positive rate. In contrast, although a few true causal QTL were missed in the preselection stage, SNPs with null effects that were moved forward into stage two were estimated to have very small effects by both ST-BayesC and MegaBayesC.

**Figure 6.**
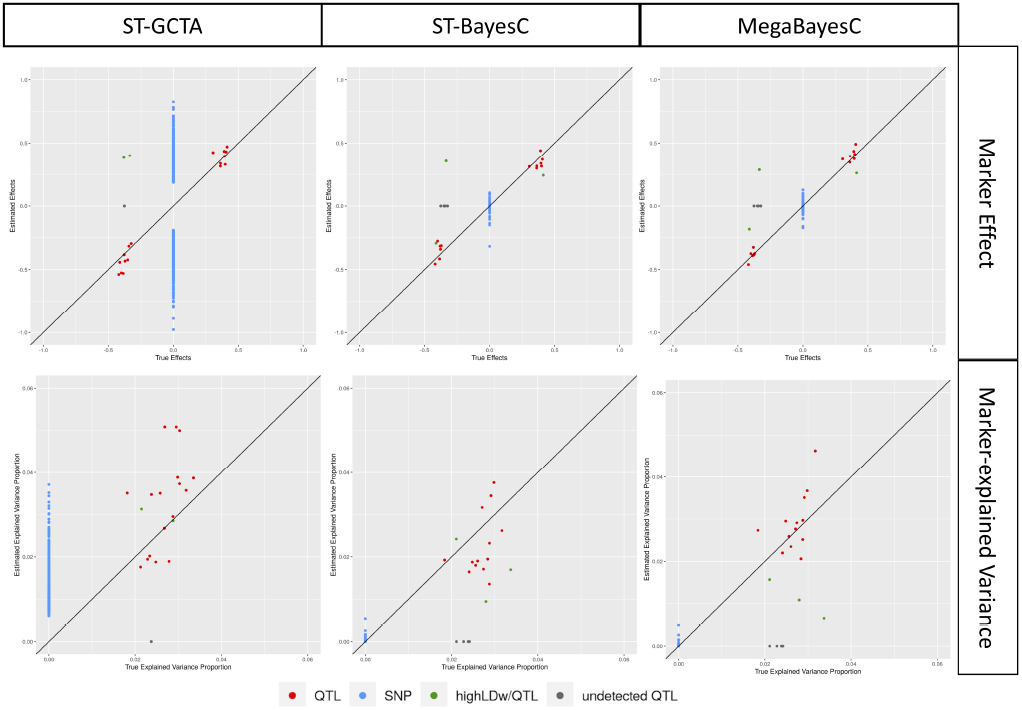
The relationship between estimated and true values of marker effects and marker-explained proportion of variance for focal trait. Three different methods (ST-GCTA, ST-BayesC, and MegaBayesC) were compared. Details of each method are presented in **Materials and Methods**.

Note that in some cases, SNPs that are in LD with true QTL were selected instead of the causal QTL. When the linkage phase was negative, the estimated effect sizes for linked SNPs have the opposite sign, which increases the reported RMSE. However, even in these cases, the proportion of variance explained by these linked markers is close to the proportion that would have been explained by the true QTL, so the effect of LD on the RMSE of marker-explained variances is minimized.

### Identifying Candidate Genes for Flowering Time in Arabidopsis using Gene Expression Data as Secondary Traits

We applied the two-stage MegaBayesC and the one-stage single-trait GWAS (ST-GCTA) to the task of identifying candidate genes that regulate flowering time in *Arabidopsis thaliana* using actual flowering time measurements and genotype data from 1003 *A. thaliana* accessions. In MegaBayesC, we included the expression of 20843 genes measured on 649 of the accessions as secondary traits.

Potentially important SNPs with marker-explained variance greater than 0.1% in MegaBayesC and potentially important SNPs with p-value smaller than 10^*−*5^ in ST-GCTA were selected as significant SNPs. MegaBayesC was better able to select a limited number of candidate SNPs based on per-marker variance explained (Figure 7) then ST-GCTA (Figure 8) by shrinking the vast majority of SNP effects close to zero.

**Figure 7.**
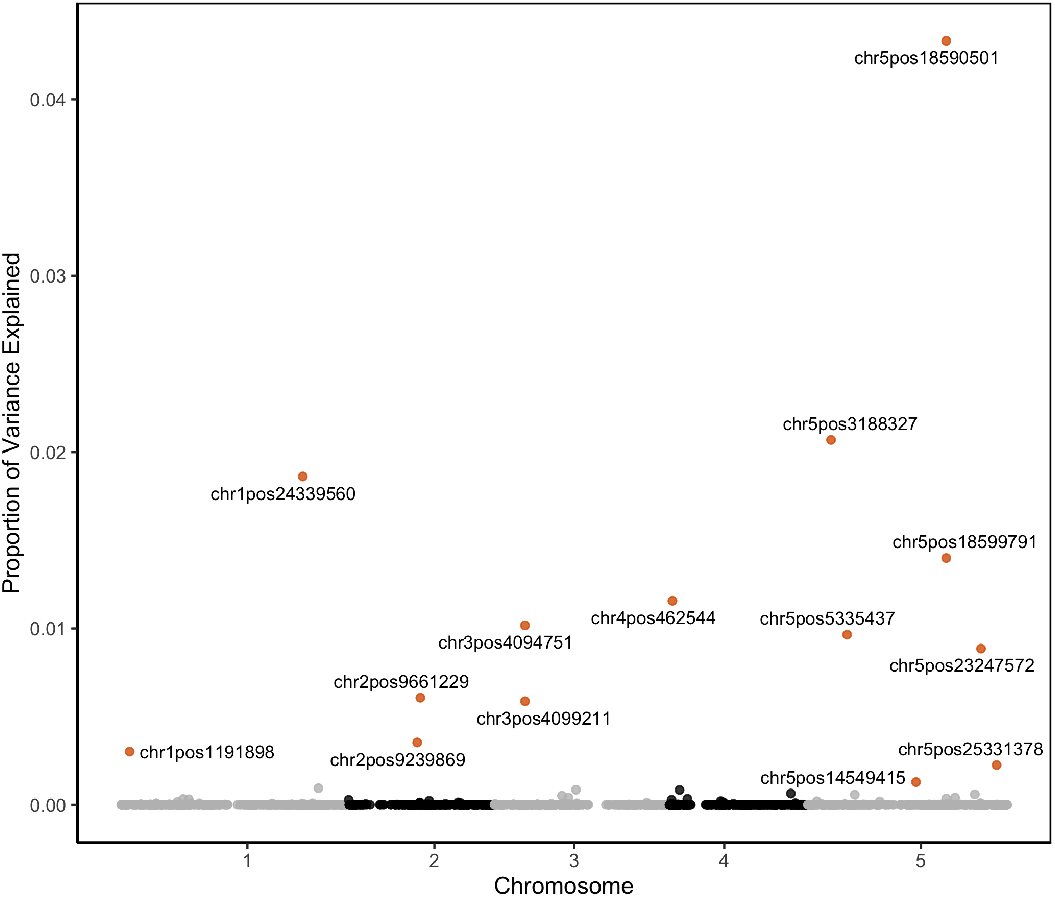
Marker-explained proportion of variance for potentially important SNPs by the two-stage analysis using MegaBayesC. The top 14 SNPs that explained the greatest proportions of variance in flowering time are highlighted.

**Figure 8.**
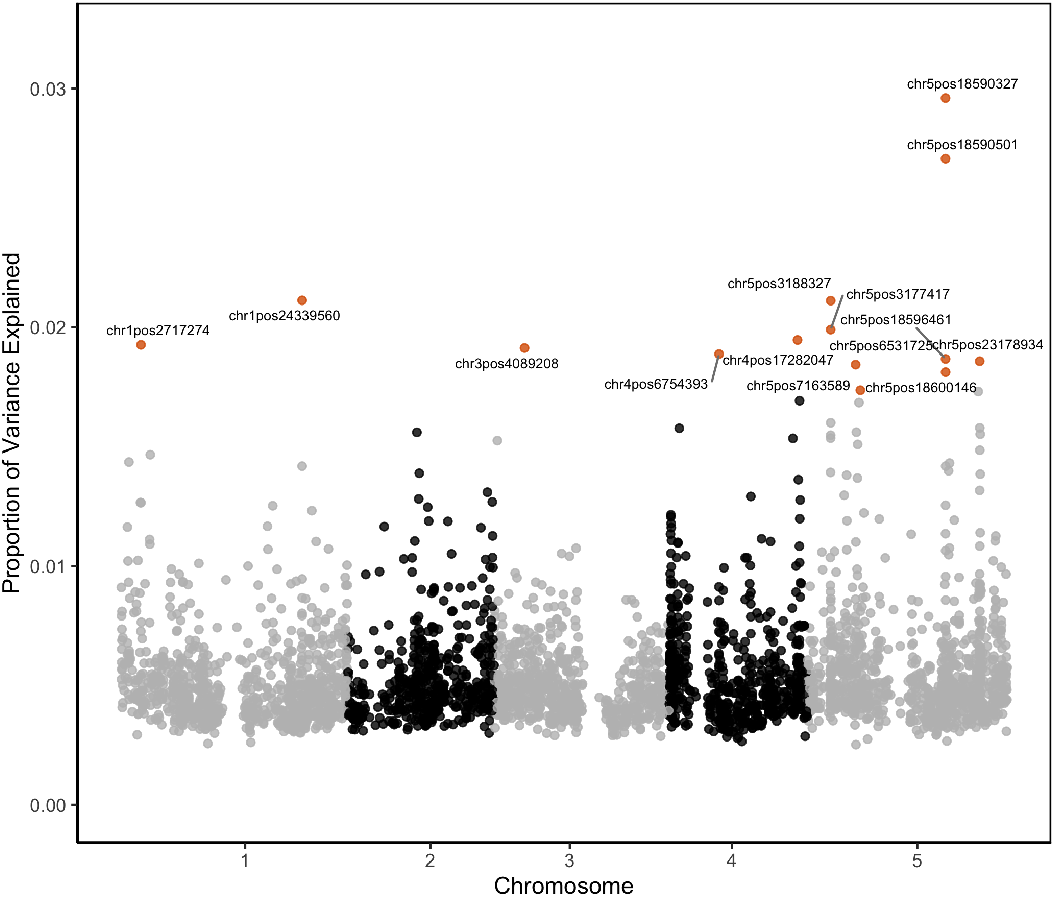
Marker-explained proportion of variance for potentially important SNPs by the one-stage ST-GCTA analysis. The top 14 SNPs that explained the greatest proportions of variance in flowering time are highlighted.

We assessed the accuracy of these associations by checking whether known flowering time-related genes are located near to the SNPs selected by each model. Using MegaBayesC, we selected 14 significant SNPs and 13 of these were located within 100Kb of known flowering time-related genes. Note that these known genes were generally not the nearest gene to the significant SNPs, but associations at this distance are not uncommon in Arabidopsis (Sasaki *et al*. 2021). For ST-GCTA, we selected 34 significant SNPs, among which 26 SNPs were located within 100Kb of known flowering time-related genes. In total, based on our prior knowledge, 14 and 15 genes were detected by MegaBayesC and ST-GCTA, respectively. Detailed comparison on detected genes between MegaBayesC and ST-GCTA is shown in Table 1.

**Table 1.**
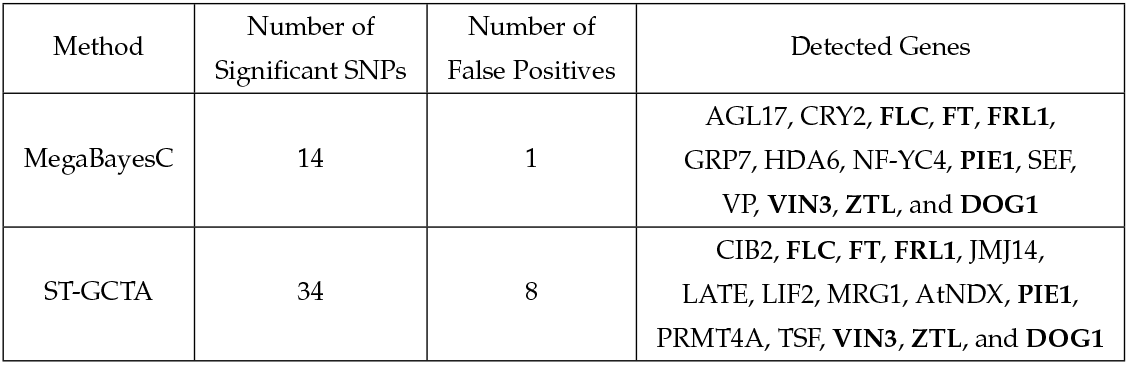
Detailed information on detected genes from ST-GCTA and MegaBayesC. Bold fonts are used to indicate genes that are detected in both methods.

## Discussion

The emergence of new types of phenotype data, such as gene expression or spectral reflectances, has created a demand for the development of robust models that are able to analyze large numbers of phenotypes in genome-enabled analysis. Although Bayesian regression models with mixture priors allow for more biologically meaningful prior assumptions on the effect size distributions of causal variants, their corresponding multivariate models (Cheng *et al*. 2018b) suffer from a high computational burden. In this paper, we developed a Bayesian sparse factor model with mixture priors on marker effects (MegaBayesianAlphabet) to implement both genome-wide prediction and association for analyses with hundreds to tens-of-thousands of phenotypes. MegaBayesianAlphabet uses a moderate number of latent factors (*K*) to account for the covariance among the observed traits. This substantially reduces the computational burden relative to either a multivariate Bayesian regression model or a multivariate linear mixed model with fully-parameterized trait covariance matrices when the number of traits (*t*) is large.

However, the sparse factor structure of MegaBayesianAlphabet does not reduce the model complexity enough to enable mixture priors over the millions of genetic markers that are available in many systems from high-density genotyping arrays or whole genome sequencing. When marker effects of the factors and the trait-specific residuals are both included in the model, the number of marker effects to be estimated is equivalent to (*t* + *K*) *×p*, with t being the number of observed traits, K being the number of factors, and p being the number of total SNPs, which would require a tremendous amount of computational time and memory storage for whole-genome analysis.

We therefore developed two approximations to greatly reduce the time complexity of the full model. First, we forced the marker effects to affect the secondary traits through the *K* factors (although we do allow marker effects to independently control the focal trait). This reduces the number of marker effects to (*K* + 1)*p*. Second, we developed a two-stage approach to prune the candidate markers before subjecting the pruned markers to the MegaBayesC analysis. For our MegaBayesC analysis of the *Arabidopsis* dataset with *n* = 649, *t* = 20844, and *p* = 2804, it took around 3 hours to sample a MCMC chain of 10,000 iterations on a computer with 1 node and 20 CPU.

While MegaBayesC, and MegaBayesianAlphabet more generally, shows promise in its ability to integrate thousands of traits in genome-wide prediction and association, the trade-off between the benefit of incorporating secondary traits and the computational cost brought from the increased model complexity must be considered. Based on our simulated study, MegaBayesC can effectively disentangle the genetic and non-genetic sources of covariation among observed traits. When there is an important environmental component in the variation of focal trait, and this environmental component is shared by many other highly correlated traits, we expect MegaBayesianAlphabet models to provide a large benefit by providing a tool to effectively control for this environmental variation. However, when the secondary traits are not highly correlated with the focal trait, or the heritability of the focal trait is already sufficiently high, MegaBayesianAlphabet may prove less useful.

In this paper, we have focused on two versions of MegaBayesianAlphabet: MegaBayesC with the BayesC prior on the marker effects, and MegaRRBLUP with a ridge prior on the marker effects. Implementing other mixture priors in the `MegaLMM` R package is relatively straightforward, and we anticipate that the BayesA, BayesB or BayesR priors may provide benefits in specific datasets.

## Appendix

### Gibbs Sampler Updates

#### Sample F given all other parameters

To sample **F**, we transpose Eq. 1:

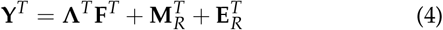

where **M**_*R*_ = **X**_1_ **B**_1_ + **X**_2*R*_ **B**_2*R*_. Conditioning on **B**_2*F*_, **B**_2*R*_, columns of **F**_*T*_ and 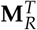 are uncorrelated and we can represent Eq. 4 as a set of simple linear regressions:

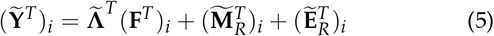

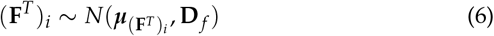

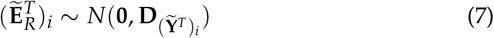

where 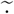 denotes the removal of missing trait data from the corresponding entity. For example, (**Y**)_*i*_ is the sub-vector of nonmissing traits in the *i*th row of 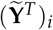 denotes the *i*th row of **F**, which follows a multivariate normal distribution with mean 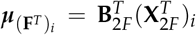 and (co)variance matrix 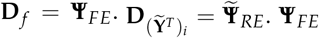 and 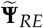 are diagonal matrices.

Let 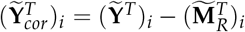, we have

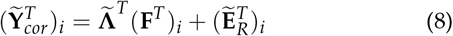

For simplicity, let 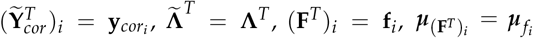 and 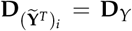. The full conditional posterior distribution for (**F**^*T*^)_*i*_ is derived as:

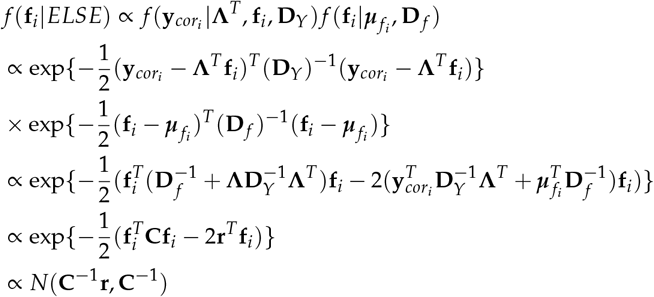

Therefore, (**F**^*T*^)_*i*_|*ELSE ∼ N*(***µ*, Σ**) with

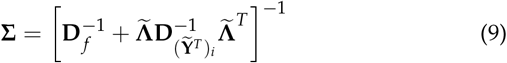

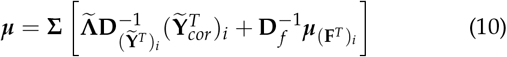

#### Sample parameters in Λ

##### *Full conditional posterior distribution of* Λ

The prior for *λ*_*j*_ is specified as follows:

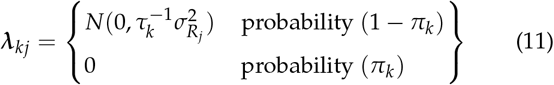

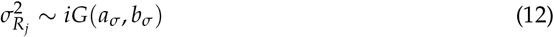

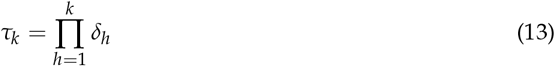

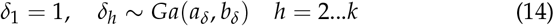

This mixture prior for ***λ***_*j*_ can be parameterized as: 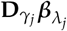, where 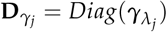with

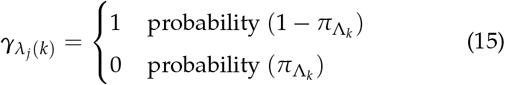

and 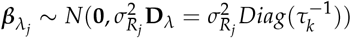 for *k* = 1, 2, … *K*.

Conditional on **F**, Eq. 1 can be simplified into t independent univariate linear mixed models for the columns of **Y**. For the *j*th column of **Y**:

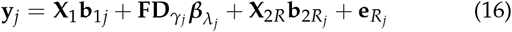

where 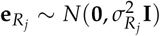.

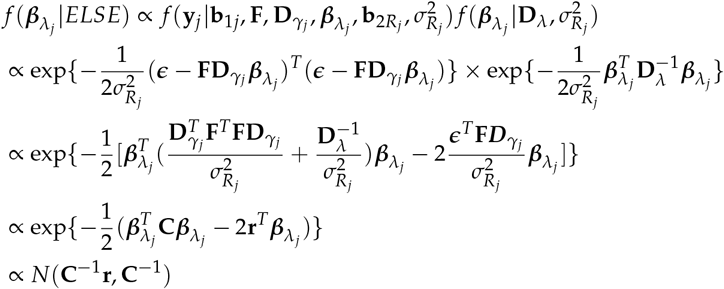

where 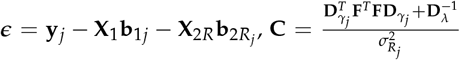, and 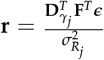.

Besides the full conditional posterior distribution for the multivariate 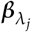 as derived above, a univariate version for the elements in 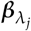 is also derived as follows to prepare for the derivation of 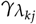.

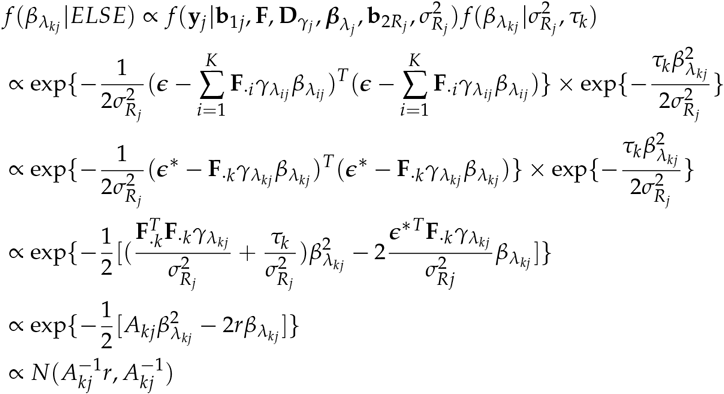

where 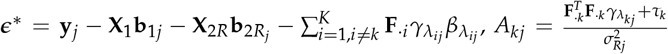, and 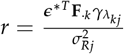.

##### Full conditional posterior distribution of 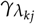

From the model specification, ***γ*** variables can take either 0 or 1. Let ***θ*** denote all other parameters except for 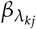and 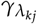, the marginal full conditional distribution of 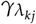 that integrates 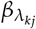is shown as:

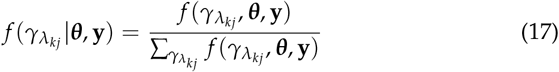

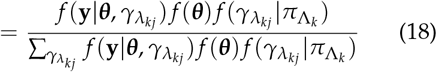

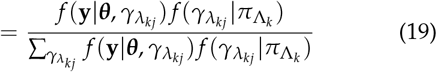

Since 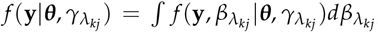, the derivation for 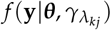is shown as follows.

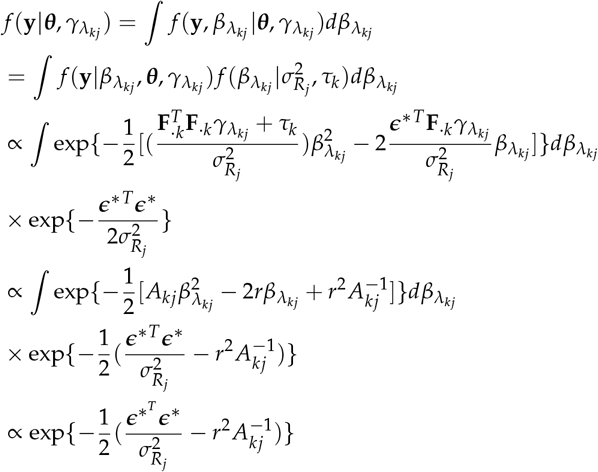

where 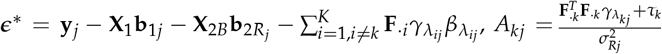, and 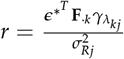.

Given Eq. 19, we have

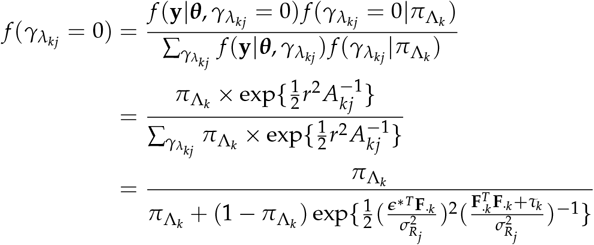

##### Full conditional posterior distribution of *δ*_*l*_

In order to sample *τ*_*k*_, we need to firstly sample *δ*_*l*_ when *K >* 1. To derive the full conditional posterior distribution of *δ*_*l*_, vectorize **Λ** as ***λ***. Then, we have

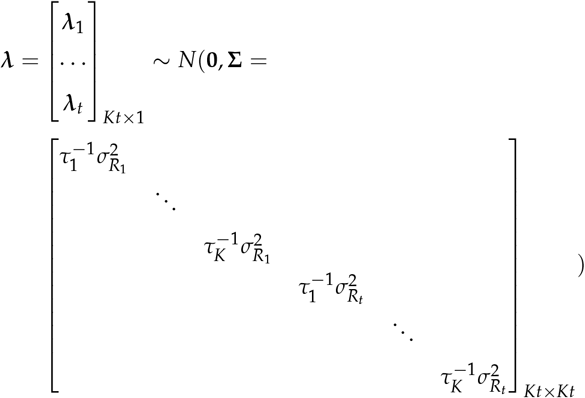

Note that the determinant of a diagonal matrix is the product of elements of its diagonal.

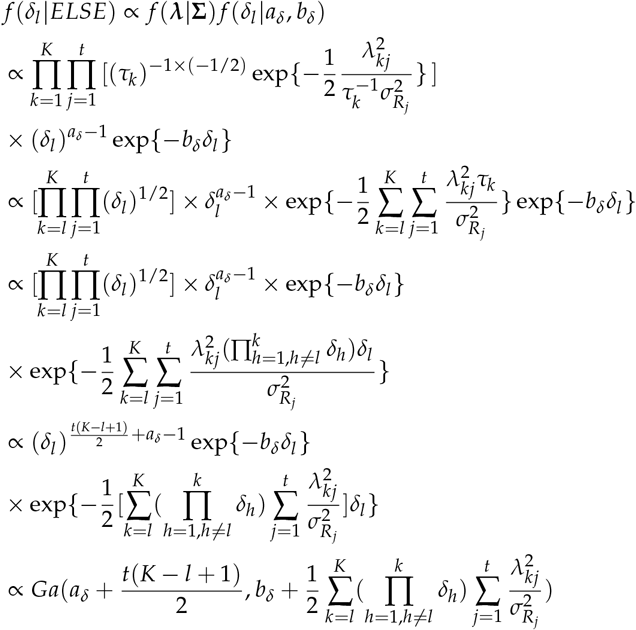

#### Parallel Model Setting

Given **F** and **Λ**, although the design matrices may differ for columns of **Y** and **F**, the form of both sets of conditional model can be similarly expressed as:

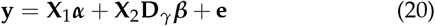

where

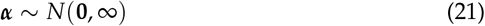

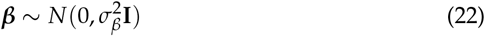

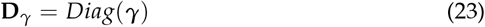

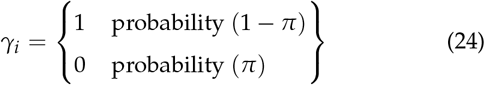

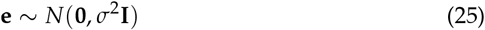

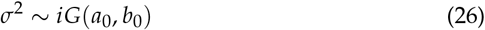

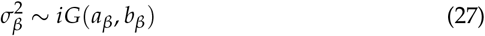

Conditional on **F** and **Λ**, Eq. 1 can be simplified into t independent univariate linear mixed models for the columns of **Y**_*cor*_ = **Y** *−* **FΛ**:

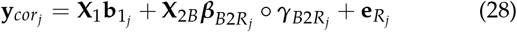

where

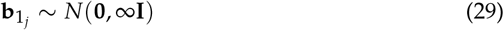

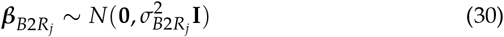

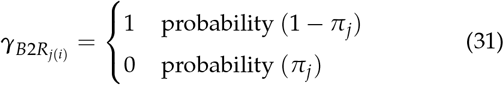

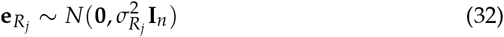

Besides the columns of **Y**, the columns of **F** (Eq. 2) can be similarly expressed into K independent univariate linear mixed models:

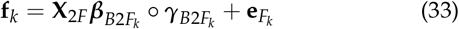

where

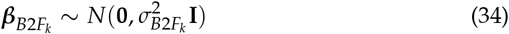

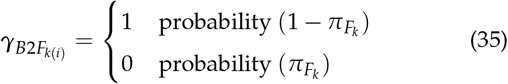

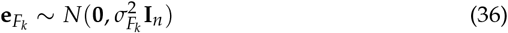

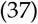

Here, factor-specific and trait-specific prior on the marker exclusion probability (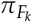and *π*_*j*_) and the variance of marker effects (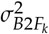and 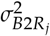) are used for each latent factor and observed trait. We can see that the columns of **Y** and **F** can be generally expressed by Eq. 20. That is, for columns of 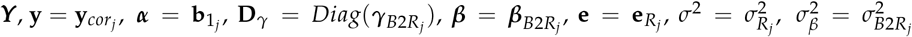. Similarly, for columns of **F, y** = **f**_*k*_, ***α*** is empty, 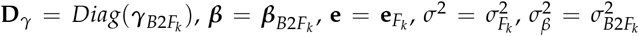. Furthermore, we defined the following term based on the notation in Eq. 20:

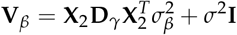

##### Full conditional posterior distribution of α

The conditional posterior distribution for ***α***(*i*.*e*., **b**_1*j*_) is derived as (integrating out ***β***):

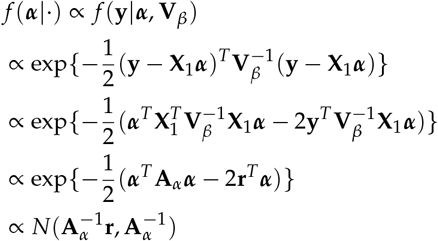

where 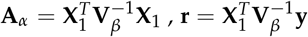. The dimension of **A**_*α*_ is *a*(*b*_1_) *× a*(*b*_1_), and the dimension of **V**_*β*_ is *n × n*.

##### Full conditional posterior distribution of *σ*^2^

The conditional posterior distribution for *σ*_2_(*i*.*e*., 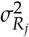 and 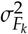) is derived as:

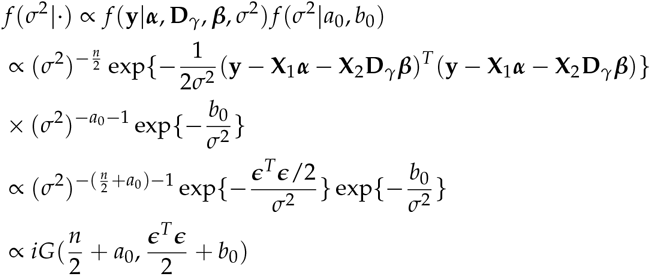

where ***ϵ = y − X***_***1***_***α − X***_***2***_***D***_***γ***_ ***β***

##### Full conditional posterior distribution of β

The conditional pos-terior distribution for ***β*** is derived as:

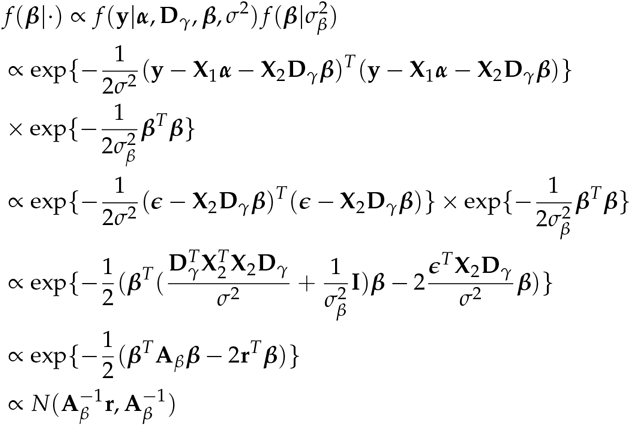

where 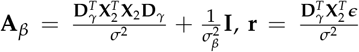. The dimension of **A**_*β*_ is *b × b*. For columns of 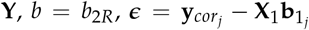. For columns of **F**, *b* = *b*_2*F*_, ***e*** = **f**_*k*_. Besides the full conditional posterior distribution of the multivariate ***β*** as derived above, a univariate version for the elements *β*_*l*_ in ***β*** is also written as follows.

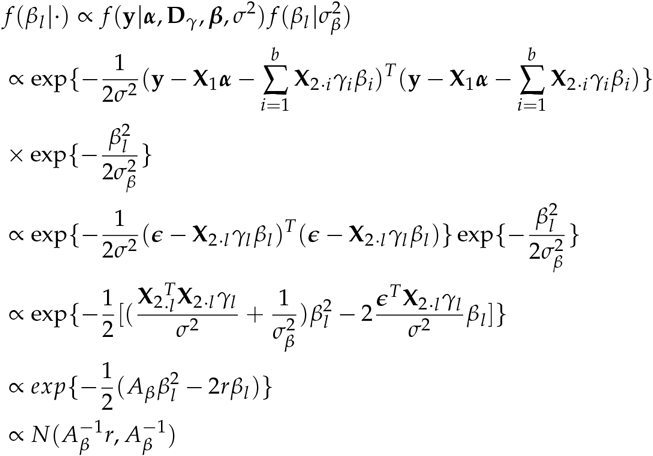

where 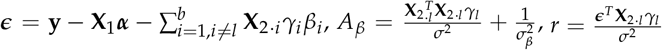.

##### Full conditional posterior distribution of *γ*_*l*_

Let ***θ*** denote all other parameters except for *β*_*l*_ and *γ*_*l*_, the marginal full con-ditional distribution of *γ*_*l*_ that integrates out *β*_*l*_ is shown as:

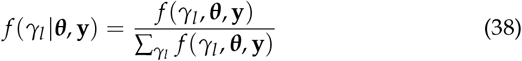

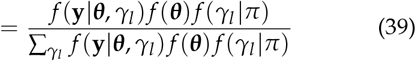

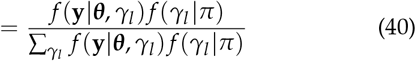

Since *f* (**y**|***θ***, *γ*_*l*_) = *f* (**y**, *β*_*l*_|***θ***, *γ*_*l*_)*dβ*_*l*_, the derivation for *f* (**y**|***θ***, *γ*_*l*_) is shown as follows.

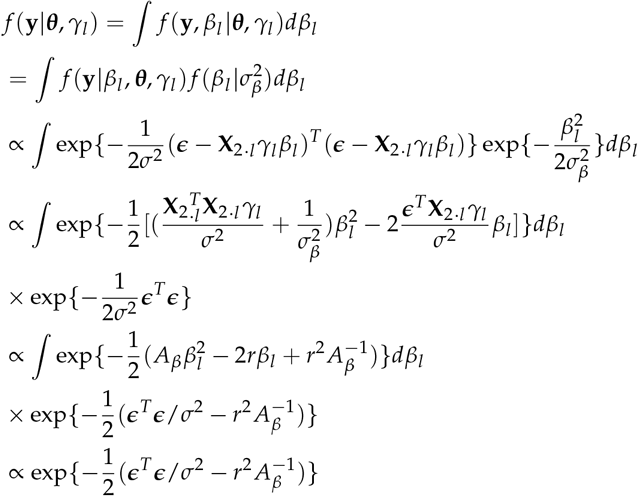

where 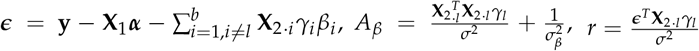.

Given Eq. 40, we have

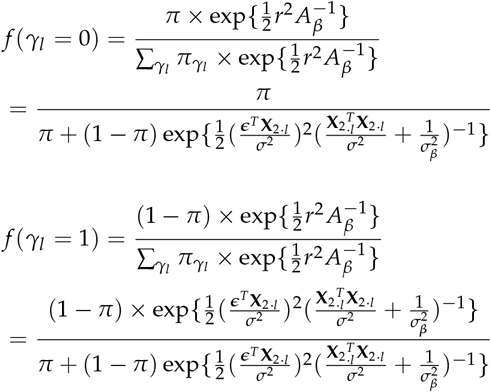

##### Full conditional posterior distribution of 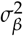

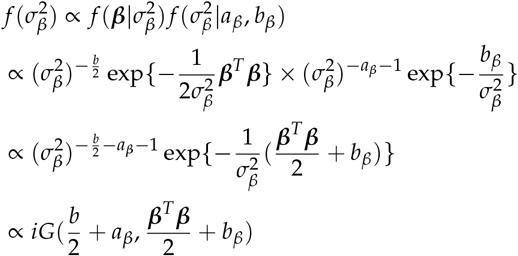

##### Specification of parameters for the real data analysis performed in the paper

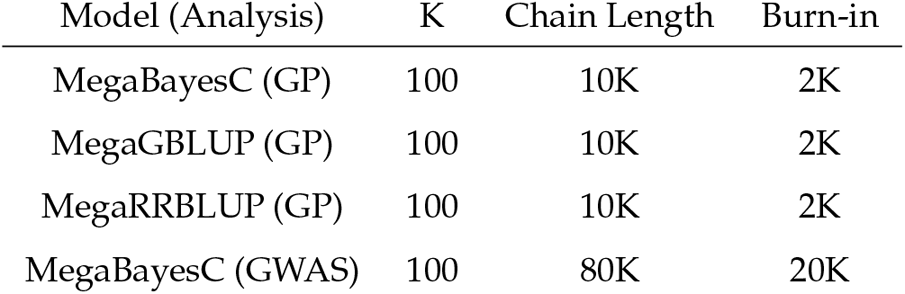

###### Supplementary Plots

**Figure 9.**
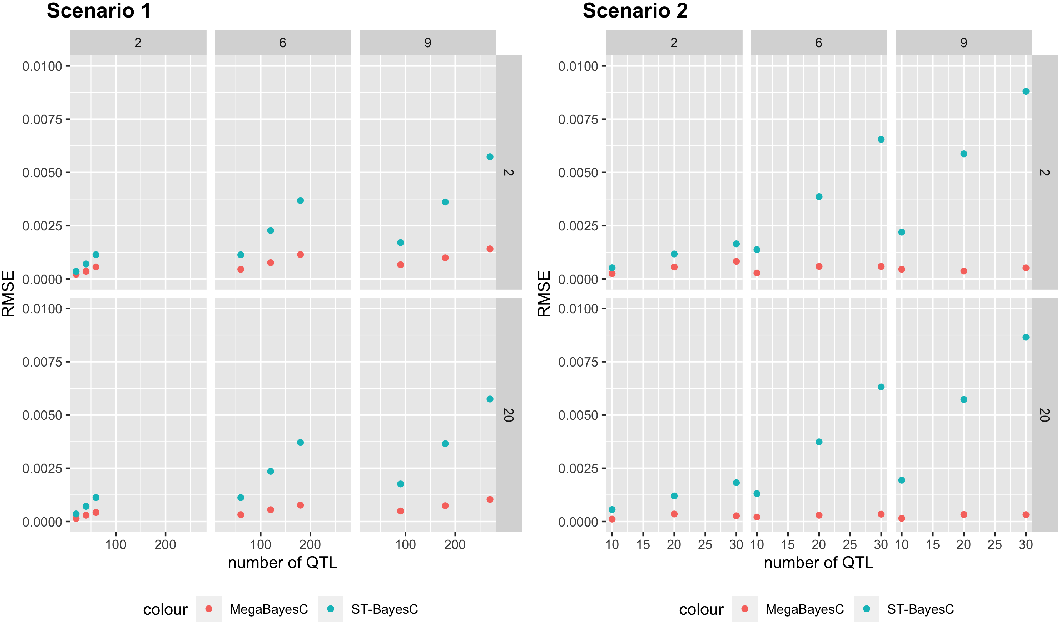
RMSE of estimated marker effects under two scenarios for the total 36 different simulation settings. The performance of single-trait BayesC and MegaBayesC were compared. The performance of models for the simulation setting with *n*_*trait*_ = 2 and 20 are presented at the first and second row, respectively. The performance of models for the simulation setting with *n* _*f actor*_ = 2, 6, 9 are presented at the first, second, and third column, respectively.

